# Small Molecule Screen Identifies Pyrimethamine as an Inhibitor of NRF2-driven Esophageal Hyperplasia

**DOI:** 10.1101/2022.12.05.519147

**Authors:** Chorlada Paiboonrungruang, Zhaohui Xiong, David Lamson, Yahui Li, Brittany Bowman, Julius Chembo, Caizhi Huang, Jianying Li, Eric W. Livingston, Jon E. Frank, Vivian Chen, Yong Li, Bernard Weissman, Hong Yuan, Kevin P. Williams, M. Ben Major, Xiaoxin Chen

**Author notes:** **Co-Corresponding authors:** Kevin P. Williams, PhD, Department of Pharmaceutical Sciences and Biomanufacturing Research Institute and Technology Enterprise, North Carolina Central University, Durham, NC 27707, USA. Michael Ben Major, PhD, Department of Cell Biology and Physiology, Washington University in St. Louis, St. Louis, MO 63110. Xiaoxin Luke Chen, MD, PhD, Cancer Research Program, Julius L. Chambers Biomedical Biotechnology Research Institute, North Carolina Central University, 700 George Street, Durham, NC 27707, USA. Equal contribution to this research work (Chorlada Paiboonrungruang, Zhaohui Xiong).

## Abstract

**Objective:** NRF2 is a master transcription factor that regulates the stress response. NRF2 is frequently mutated and activated in human esophageal squamous cell carcinoma (ESCC), which drives resistance to chemotherapy and radiation therapy. Therefore, a great need exists for NRF2 inhibitors for targeted therapy of NRF2^high^ ESCC.

**Design:** We performed high-throughput screening of two compound libraries from which hit compounds were further validated in human ESCC cells and a genetically modified mouse model. The mechanism of action of one compound was explored by biochemical assays.

**Results:** Using high-throughput screening of two small molecule compound libraries, we identified 11 hit compounds as potential NRF2 inhibitors with minimal cytotoxicity at specified concentrations. We then validated two of these compounds, pyrimethamine and mitoxantrone, by demonstrating their dose- and time-dependent inhibitory effects on the expression of NRF2 and its target genes in two *NRF2*^*Mut*^ human ESCC cells (KYSE70 and KYSE180). RNAseq and qPCR confirmed the suppression of global NRF2 signaling by these two compounds. Mechanistically, pyrimethamine reduced NRF2 half-life by promoting NRF2 ubiquitination and degradation in KYSE70 and KYSE180 cells. Expression of an *Nrf2*^*E79Q*^ allele in mouse esophageal epithelium (*Sox2CreER;LSL-Nrf2*^*E79Q/+*^) resulted in an NRF2^high^ phenotype, which included squamous hyperplasia, hyperkeratinization, and hyperactive glycolysis. Treatment with pyrimethamine (30mg/kg/day, *p*.*o*.) suppressed the NRF2^high^ esophageal phenotype with no observed toxicity.

**Conclusion:** We have identified and validated pyrimethamine as an NRF2 inhibitor that may be rapidly tested in the clinic as a radiation and chemotherapy sensitizer for NRF2^high^ ESCC.

**Summary:** *What is already known on this topic – summarise the state of scientific knowledge on this subject before you did your study and why this study needed to be done:*

- Mutational activation of the NRF2 transcription factor drives ESCC progression and therapeutic resistance. Targeted therapies to block NRF2 have not yet been realized, despite great needs.

*What this study adds – summarise what we now know as a result of this study that we did not know before:*

- A screen of >35,000 small molecules identified eleven potential NRF2 inhibitors. Pyrimethamine and mitoxantrone were validated to inhibit the expression of NRF2 and NQO1 in human ESCC cells in both dose- and time-dependent manners.
- Pyrimethamine enhanced NRF2 protein ubiquitination and degradation, resulting a decreased half-life.
- A genetically modified mouse model was established to express the *Nrf2*^*E79Q*^ mutant allele in the mouse esophageal epithelium upon tamoxifen induction. Pyrimethamine suppressed the NRF2^high^ esophageal phenotype induced by the mutant allele.

*How this study might affect research, practice or policy – summarise the implications of this study:*

- As an FDA-approved drug, Pyrimethamine has the potential for immediate translation to a clinical trial on NRF2^high^ ESCC in humans.
- Further exploration of its mechanisms of action may lead to more potent NRF2 inhibitors for future use.

## Introduction

The development of human esophageal squamous cell carcinoma (ESCC) evolves through histological stages of hyperplasia, dysplasia, and carcinoma. The 5-year survival rate for ESCC is ∼18%, a number that reflects late diagnosis, the aggressiveness of the disease, and a lack of effective treatment strategies^1, 2^. Thus, there is a great need to further elucidate the molecular mechanisms driving the etiology of ESCC and develop more effective treatment strategies.

Nuclear factor (erythroid-derived 2)-like 2 (*NRF2* or *NFE2L2*) mutations are commonly seen in ESCC, with frequencies of 10-22% ^3, 4^. Mutations in other genes of the NRF2 signaling pathway, Kelch-like ECH-associated protein 1 (*KEAP1*) and Cullin 3 (*CUL3*), are present but less common than those in *NRF2. NRF2* mutations mainly occur in hotspots localized to the DLG and ETGE motifs, two domains required for NRF2 association with KEAP1, its primary inhibitor ^5^. The NRF2 signaling pathway is recognized as a double-edged sword in the context of carcinogenesis^6, 7^. On one hand, chemical or genetic activation of NRF2 induces cytoprotective enzymes conferring protection against chemical carcinogenesis in multiple models including esophageal cancer ^8, 9^. On the other hand, NRF2 activation mitigates stress associated with onco-metabolism, hypoxia, immune pressure, and aberrant proliferation. Indeed, NRF2 hyperactivation governs many of the cancer hallmarks, including cell proliferation, differentiation, immune infiltration, and cancer metabolism^10^. NRF2^high^ cancers of varied tissue origins demonstrate resistance to immune checkpoint inhibitors, chemotherapy, and radiation therapy ^11, 12^. In human ESCC, NRF2 overexpression significantly correlates with increased lymph node metastasis, postoperative recurrence, and decreased overall survival ^13-16^. Although not yet realized, the NRF2 signaling pathway is regarded as a tractable molecular target for cancer therapy^17^. In mice, NRF2 hyperactivation in *Keap1*^*-/-*^ mice results in esophageal hyperplasia and hyperkeratosis^18^. NRF2 is a critically important KEAP1 substrate as the esophageal phenotype of *Keap1*^*-/-*^ mice is rescued in *Nrf2*^*-/-*^*;Keap1*^*-/-*^ and *K5Cre;Nrf2*^*fl/fl*^*;Keap1*^*-/-*^ mice^18, 19^.

Approximately 30 small-molecule NRF2 inhibitors have been reported^20, 21^. While invaluable as tool compounds, these inhibitors have yet to be clinically proven. Continued and enhanced drug development efforts that identify potent, specific, and efficacious drugs for NRF2^high^ cancer are greatly needed. In this study, we used high-throughput small molecule screens to identify NRF2 inhibitors. We present data validating two of these drugs, including mechanistic insights and *in vivo* efficacy in an NRF2^high^ genetically engineered mouse model (GEMM) of esophageal hyperplasia.

## Materials and Methods

### Cell culture and chemicals

*NQO1-YFP* H1299 cells (human lung adenocarcinoma), and NRF2^high^ KYSE70 (*NRF2*^*W24C*^) and KYSE180 (*NRF2*^*D77V*^) and NRF2^low^ KYSE450 (*NRF2*^*WT*^) cells (human ESCC-DSMZ, Braunschweig, Germany) ^20^ were cultured in Gibco RPMI1640 GlutaMAX (ThermoFisher, Waltham, MA) supplemented with 10% FBS and 1% antibiotics (penicillin/streptomycin). *NQO1-YFP* H1299 cells contained a YFP fragment in *NQO1*_intron 1 which responded to NRF2 activators ^22^. All cell lines were cultured in a humidified incubator at 37 °C with 5% CO_2_.

The Prestwick library (1,280 FDA-approved drugs) and Asinex library (34,560 compounds) were acquired from Prestwick Chemical (San Diego, CA) and Asinex (Winston-Salem, NC), respectively. Pyrimethamine (PYR), mitoxantrone dihydrochloride (MIT), brusatol, cycloheximide (CHX), and MG132 were purchased from Sigma-Aldrich (St. Louis, MO). All cell culture and biochemical assays were performed in duplicate or triplicate to ensure reproducibility.

### High-throughput screening

*NQO1-YFP* H1299 cells were seeded at a density of 1,000 cells/30 µl/well in 384-well plates using a Multidrop 384 (Thermo Fisher, Waltham, MA) bulk dispenser and cultured overnight. Individual compounds (1 µM in 0.1% DMSO) were added to each well using a Biomek NX workstation (Beckman-Coulter)(1 well per compound), with 0.1% DMSO as the positive control and an NRF2 activator (CDDO, 200 nM in 0.1% DMSO) as the negative control. All plates were monitored using the IncuCyte S3 (Essen BioScience, Ann Arbor, MI). Compounds that inhibited >50% of CDDO-activated signal (green GFP signal) and allowed >80% cell survival (red mCherry signal) were further examined in a dose-dependent experiment. Individual compounds were dispensed at 10-point serial dilutions starting from 10 µM in triplicates using an HP D300 Digital Dispenser (Hewlett-Packard, Palo Alto, CA). A well-established NRF2 inhibitor, brusatol, was used as a control. The image analysis was captured at 0, 24, 48, and 64 h. GraphPad Prism was used to show the dose-dependent screening data and calculate IC_50_.

Cytotoxicity of 12 compounds (8 Prestwick compounds and 4 Asinex compounds) was tested in a dose-dependent screening assay. KYSE70 and KYSE180 cells were seeded at a density of 1,200 cells/30µl/well in 384-well plates and cultured overnight. Compounds were added to the plate at 2-point serial dilution in triplicate starting from 10 µM with 0.1% DMSO as the negative control and 30 μM of benzethonium chloride as the positive control. After 3 days, another positive control was treated with 0.1% Triton X-100 right before staining. The viability of each well was assessed by double fluorescent staining with 0.5 μg/ml Hoechst and 100 nM YOYO1 (Invitrogen, Waltham, MA). Four images for each well were captured under 4X magnification using Thermo Scientific CellInsight NXT high-content screening platform. GraphPad Prism was applied to calculate the normalized percentage of cell count and the normalized percentage of dead cells. IC_50_ was calculated and graphs were generated for 11 compounds. PYR (an FDA-approved antimalarial and anti-toxoplasmosis drug targeting dihydrofolate reductase, DHFR) ^23^ and MIT (an FDA-approved chemotherapeutic drug) ^24^ were chosen for further validation because of their relatively potent NRF2-inhibitory activities and weak cytotoxic activities at the concentrations studied (Table 1).

**Table 1.** Effects of 11 hit compounds on cell viability and NRF2 activity in H1299-YFP cells, and cell proliferation and cell death in NRF2^high^-ESCC cells (KYSE70 and KYSE180)

| Compound | H1299-NQO1-YFP |  | KYSE70 |  | KYSE180 |  |
| --- | --- | --- | --- | --- | --- | --- |
|  | Cell viability | NRF2 inhibition | Cell proliferation | Cell death | Cell proliferation | Cell death |
| | (IC<sub>50</sub>, μM) | (IC<sub>50</sub>, μM) | (IC<sub>50</sub>, μM) | (IC<sub>50</sub>, μM) | (IC<sub>50</sub>, μM) | (IC<sub>50</sub>, μM) |
| Isoproterenol hydrochloride | >10 | 0.84 | 5.2 | >10 | >10 | >10 |
| Ethoxyquin | >10 | 0.29 | >10 | >10 | >10 | >10 |
| Mitoxantrone dihydrochloride (MIT) | >10 | 0.035 | 5.84 | 3.24 | 4.57 | 5.22 |
| Isoetharine mesylate | >10 | 0.49 | >10 | >10 | >10 | >10 |
| Isoproterenol bitrtrate | >10 | 0.43 | 4.9 | >10 | >10 | >10 |
| Pyrimethamine (PYR) | 1.17 | 0.23 | >10 | 14.6 | 13.7 | >10 |
| Triamterene | 3.70 | 0.72 | 7.7 | >10 | >10 | >10 |
| Eseroline fumarate | >10 | 0.26 | 10 | >10 | >10 | >10 |
| Asinex 1 | 2.99 | 0.47 | 2.8 | >10 | 3.2 | >10 |
| Asinex 2 | 4.87 | 0.04 | 6.3 | >10 | 15.3 | >10 |
| Asinex 3 | >10 | 1.21 | 5 | >10 | >10 | >10 |

### Western blotting and immunoprecipitation

The nuclear and cytosolic extracts were prepared using a CelLytic NuCLEA extraction kit (Millipore Sigma) or total protein (whole cell lysate) using RIPA buffer. Proteins were separated by SDS-PAGE and transferred to nitrocellulose membranes. Membranes were blocked and then incubated with a primary antibody overnight at 4°C (Table S1). Chemiluminescence was detected using autoradiography film and followed by quantification using ImageJ.

To analyze NRF2 ubiquitination, cells were treated with a compound and then treated with or without MG132 (10 µM) before being harvested in RIPA buffer. The cell lysate was pre-cleaned with protein G agarose beads (Roche Diagnostic, Basel, Switzerland) for 1 h at 4°C. Samples were incubated with 1 µg anti-NRF2 antibody overnight at 4°C with rotation, and then 25 µl protein G agarose beads were added for 1.5 h at 4°C. Immunoprecipitants were washed and boiled in 2×SDS loading buffer at 95 °C for 5 min, and analyzed by Western blotting with anti-ubiquitin.

### Biochemical assays

A DHFR activity assay kit (Sigma) was used to analyze the enzymatic activity. The procedure was adjusted to a reaction volume of 200 μl. The reaction progress was followed by monitoring the decrease in A_340nm_ over time. MTX (1 µM) was used as a control DHFR inhibitor.

A protein synthesis assay kit (Cayman Chemicals, Ann Arbor, MI) was used to analyze global translation according to the manufacturer’s instructions. KYSE70 cells were seeded at a density of 2000 cells/100µl/well in a 96-well plate and treated with PYR, MIT, or CHX in triplicate (positive control). Cells were examined by ImageExpress Pico (Molecular Devices, San Jose, CA).

NRF2-ARE binding assay was performed with an ELISA-based TransAM NRF2 Assay Kit (Active Motif, Carlsbad, CA) according to the manufacturer’s instructions. Briefly, nuclear extracts were prepared from KYSE70 cells and added to wells (20 μg/well) containing the ARE oligonucleotide. After incubation for 1 h with mild agitation and proper washing, anti-NRF2 was added and incubated for 1 h. An HRP-conjugated secondary antibody and a substrate were added sequentially for measuring A_450nm_.

NRF2 half-life was analyzed with a CHX chase analysis. Cells were pre-incubated with PYR (10 μM), MIT (20 nM), or MTX (50 nM) for 4 h, followed by the addition of 200 μg/ml CHX was to block protein synthesis. The cells were collected at the specified time points after CHX treatment. Total protein lysate was separated by SDS-PAGE and blotted with antibodies against NRF2 and GAPDH. Western blot images were quantified with ImageJ for the calculation of NRF2 half-life.

RNAseq was performed by Novogene (Durham, NC). A total amount of 0.4 µg RNA per sample was used as input material for the RNA sample preparations. Sequencing libraries were generated using NEBNext Ultra TM RNA Library Prep Kit for Illumina (NEB, Ipswich, MA) following the manufacturer’s recommendations and index codes were added to attribute sequences to each sample. After library preparation, the samples were sequenced (150 bp) on Novaseq 6000 according to manufacturer specifications. The quality-filtered reads were aligned with STAR (version 2.6.90c) to the human reference genome (hg38) with its respective RefSeq annotation, and the expression levels of genes were obtained with FeatureCounts (version 1.5.1). DESeq2 (version 1.34.1) was used to identify the differentially expressed genes (DEGs). Data matrices were normalized and the DEGs were reported according to the fold change cut-off and corrected modeling p-values. Gene set analysis (GSA) was done to evaluate the differential enrichment of transcription factor (TF) gene sets, knowledge-based (KB) gene sets, gene ontology (GO) gene sets, and canonical pathway (CP) gene sets. The raw data has been submitted to the NCBI GEO database (GSE173164). qPCR was performed to quantify the expression levels of genes of interest with relevant primers and TaqMan probes in a 96-well optical plate on an ABI 7900HT Fast Real-Time PCR system (Applied Biosystems).

### Animal studies

All animal experiments were approved by the IACUC at North Carolina Central University (protocol number XC06142019). *Sox2CreER* mice and *LSL-Nrf2*^*E79Q/+*^ mice^25^ were crossed to express the *Nrf2*^*E79Q*^ mutant in the esophageal epithelium of adult *Sox2CreER;LSL-Nrf2*^*E79Q/+*^ mice after tamoxifen induction (75 mg/kg/day, *i*.*p*., 5 days). When the mice were sacrificed, BrdU (50mg/kg, *i*.*p*.) was given two hours before. A segment of the esophagus was harvested and fixed in formalin for histology, while esophageal epithelium and forestomach were stored in liquid nitrogen for molecular analyses.

Six mice were assessed by ^18^F-FDG PET/CT at Week 0 (before tamoxifen induction) and Week 1 (after tamoxifen induction) following an established procedure ^26^. In brief, each mouse was orally administered with a contrast agent (Iohexol, 150 mg/ml, 0.05 ml) under light anesthesia with isoflurane for visualization of the esophagus under contrast-enhanced CT. About 350 µCi of ^18^F-FDG was injected through a tail vein catheter. PET/CT imaging was conducted at 40 min post tracer injection under anesthesia. PET images were reconstructed using 3D-OSEM method with scatter, random, attenuation, and decay correction, and registered to the CT images of the same animal. The esophagus region was manually demarcated in CT images (highlighted by Iohexol) and superimposed on PET images. The mean standardized uptake value (SUV_mean_) of ^18^F-FDG was quantified in the esophagus of each mouse.

PYR was dissolved in N-methyl-2-pyrrolidone:PEG300 at the ratio of 1:9 for *in vivo* experiments. A dose-finding experiment was carried out in adult wild-type mice to determine the proper dose of PYR (15, 30, and 60 mg/kg, *p*.*o*., 1/day for 4 weeks, n=5 per group). Since myelosuppression is a common complication of PYR treatment in humans^27^, we monitored general health (grooming, physical strength, and movement) and body weight during the treatment. Blood samples were collected for a complete blood count (IDEXX, Westbrook, ME).

Two experiments were performed to examine the therapeutic effect of PYR on the NRF2^high^ esophageal phenotype in the *Sox2CreER;LSL-Nrf2*^*E79Q/+*^ mice. In the first experiment, PYR was administered during tamoxifen induction (Figure 5A). All tissue samples were collected at Week 5. In the second experiment, PYR was administered after tamoxifen induction (Figure S6A). Part of the forestomach was harvested right after tamoxifen induction by surgical resection at Week 1, and the rest of the forestomach and esophageal epithelium were harvested when the mice were sacrificed at Week 3. Total protein was extracted from frozen tissues with a standard method for Western blotting. Paraffin sections were used for IHC of NRF2 and its target genes.

### Histochemical and immunohistochemical staining (IHC)

Formalin-fixed tissues were processed, embedded in paraffin, and serially sectioned for staining. H&E staining was conducted using a routine protocol. For IHC, antigens were retrieved on the deparaffinized sections for detection with primary antibodies (Table S1) and further detected with a streptavidin-peroxidase reaction kit and DAB as a chromogen (ABC kit; Vector Labs, Burlingame, CA). To ensure the specificity of the primary antibody, control tissue sections were incubated in the absence of the primary antibody.

### Statistical analysis

Z’ as a measure of screen assay quality was calculated according to an established method ^28^. Data were presented as the mean ± SD after the quantitation of Western bands. Most statistical analyses were performed with a two-tailed Student’s t-test. Complete blood counts were analyzed with an ANOVA test. Statistical significance was displayed as \**P*<0.05, **P<0.01, and ***P<0.001 unless indicated otherwise.

## Results

### High-throughput screening for NRF2 inhibitors

Using *NQO1*-YFP H1299 cells, we screened 35,840 chemical compounds at 1 µM (1,280 from the Prestwick library and 34,560 from the Asinex library) with the YFP green signal as an indicator of the NRF2 activity and the constitutively expressed mCherry red signal as an indicator of cell number and viability. The screen was performed at 64 h (Z’ = 0.62). Twenty-six compounds (1 µM) that inhibited >50% of the NRF2 activity and allowed at least 80% cell viability were selected for further analysis for a dose-response study using the same screening method. The list was narrowed down to 11 compounds that showed potent NRF2-inhibitory activity and low toxicity on *NQO1*-YFP H1299 cells (Figure 1 and Figure S1).

**Figure 1.**
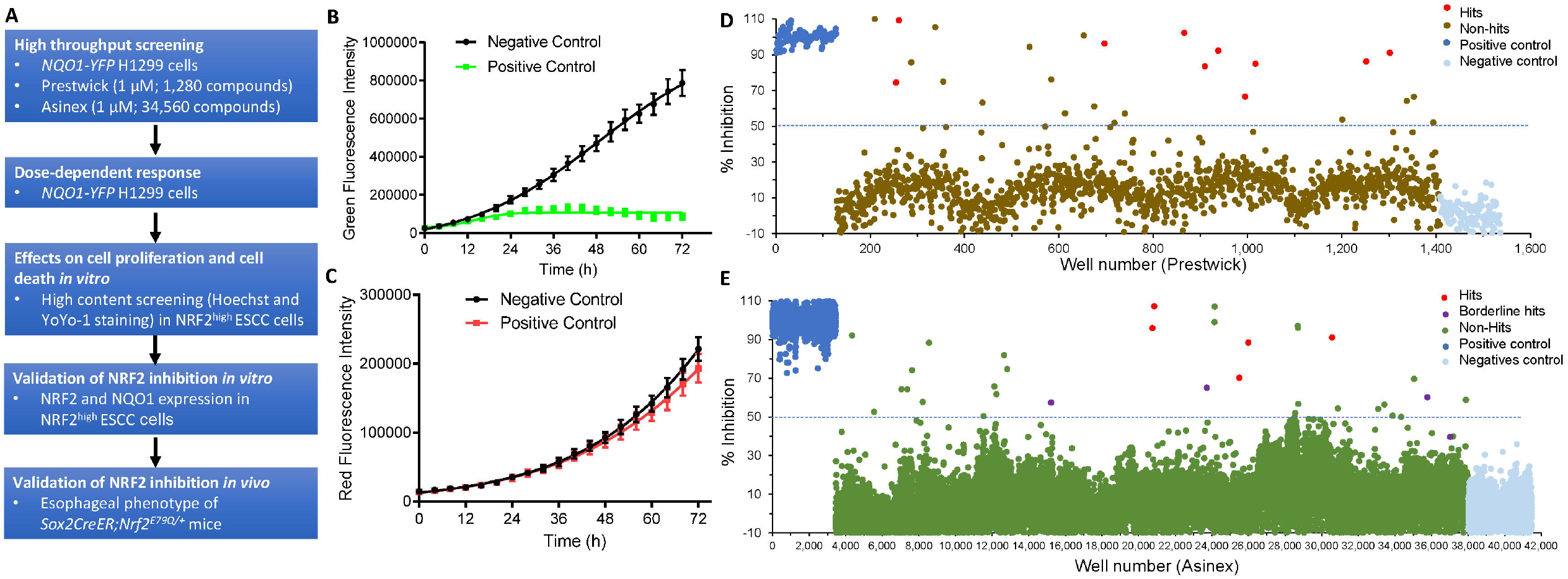
High-throughput screening for NRF2 inhibitors from two compound libraries (1 µM) using *NQO1-YFP* H1299 cells. (A) Flow chart of the screening strategy; (B) Time-dependent changes of the YPF signal (NRF2 activity); (C) Time-dependent changes of the mCherry signal (cell viability); Negative control: exposure to a known NRF2 activator (CDDO, 200 nM); Positive control: exposure to the vehicle. (D) Prestwick library (1,280 FDA-approved drug compounds); (E) Asinex library (34,560 compounds). Hit compounds were selected if the CDDO-induced NRF2 activity was inhibited by >50% and cell survival was >80%. Three compounds from the Asinex library were defined as borderline hits because their inhibition of NRF2 activity was >50%, yet their effects on cell survival were ∼80%.

We further analyzed their dose-dependent effects on the proliferation and cytotoxicity of KYSE70 and KYSE180 cells by high-content screening (Table 1, Figure S2). Two compounds, PYR (IC_50_=0.23 µM) and MIT (IC_50_=0.035 µM), were chosen for further studies due to their potent NRF2-inhibitory effects and relatively low toxicity. PYR is an FDA-approved anti-malarial and anti-toxoplasmosis drug targeting plasmodial DHFR at a concentration that is reported to be 1,000 times less than that required to inhibit the mammalian enzyme ^23^. MIT is approved by FDA for the treatment of numerous human cancers but not esophageal cancer ^24^.

### PYR and MIT inhibited NRF2 expression in *NRF2*^*Mut*^-ESCC cells *in vitro*

*NRF2*^*Mut*^-ESCC cells (KYSE70 and KYSE180) were treated with PYR and MIT to validate their inhibitory effects on NRF2 expression. Both PYR and MIT inhibited the expression of nuclear NRF2 and cytoplasmic NQO1 in a dose- and time-dependent manner in KYSE70 cells (Figure 2), as well as in KYSE180 cells (Figure S3).

**Figure 2.**
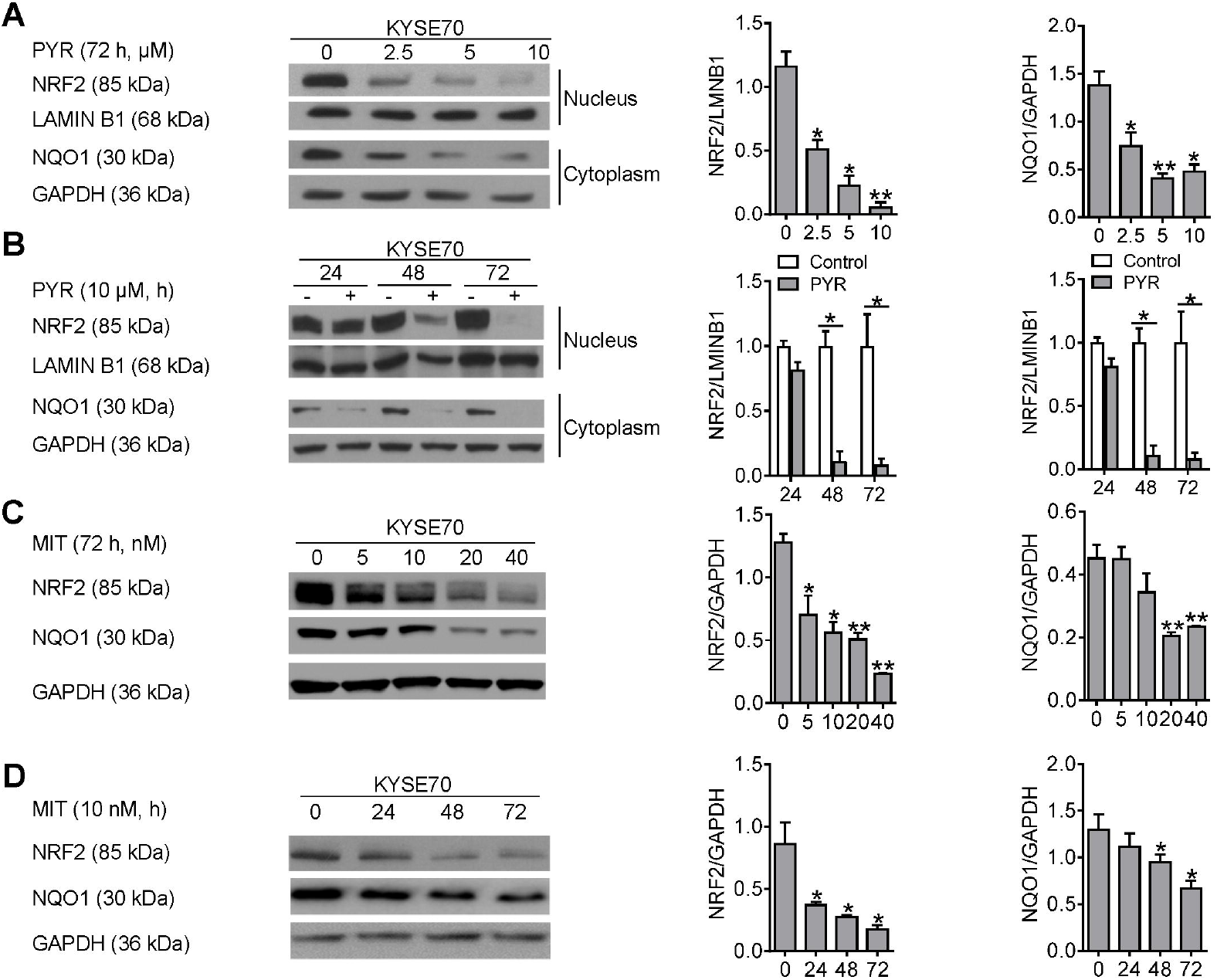
PYR and MIT downregulated NRF2 and NQO1 expression in KYSE70 cells in a dose- and time-dependent manner. (A) Dose-dependent downregulation of the expression of nuclear NRF2 and cytoplasmic NQO1 by PYR; (B) Time-dependent downregulation of the expression of nuclear NRF2 and cytoplasmic NQO1 by PYR (10 µM); (C) Dose-dependent downregulation of NRF2 expression by MIT; (D) Time-dependent downregulation of NFR2 expression by MIT (10 nM). * p<0.05, ** p<0.01

Global mRNA expression profiles by RNAseq in KYSE70 cells modified by PYR (10 µM for 72 h) and MIT (20 nM for 72 h) are shown in Figure S4A. As expected, many canonical NRF2 target genes were enriched in the control samples, as well as a gene set “differential genes in NRF2^high^ human ESCC” (Excel S1). qPCR confirmed the downregulation of *NFE2L2* mRNA (Figure S4B) and several NRF2 target genes (*AKR1B10, AKR1C2, AKR1C3, PGD, SLC7A11, TKT*) by PYR and MIT (Figure S4C). These results demonstrated that PYR and MIT inhibited NRF2 expression and transcriptional activity in *NRF2*^*Mut*^-ESCC cells *in vitro*.

### Biochemical effects of PYR and MIT on human ESCC cells *in vitro*

Since PYR was recently reported to inhibit DHFR and STAT3 in human cancer cells ^29, 30^, we evaluated whether PYR inhibited STAT3 and DHFR activity in human ESCC cells. In KYSE70 cells, PYR (up to 10 µM) did not significantly inhibit pSTAT3 and STAT3 expression (Figure S5A), but did significantly inhibit DHFR activity starting at 5 µM (Figure S5B). Although MIT was reported to inhibit topoisomerase II and the expression of an NRF2 target gene (ABCG2) in the literature^31^, MIT increased ABCG2 and TOP2A expression in KYSE70 cells (Figure S5C, S5D). Because some small molecule NRF2 inhibitors (e.g., brusatol and halofuginone) suppressed global translation^32, 33^, we next examined whether PYR and MIT inhibited global translation using a protein synthesis assay with CHX as a positive control. Neither PYR (10 µM) nor MIT (20 nM) inhibited global translation (Figure S5E). We also examined the effects of PYR or MIT on NRF2-ARE binding in KYSE70 cells and did not observe any changes (Figure S5F).

Since disruption of NRF2-KEAP1 interaction due to stress or gene mutations extends the half-life of the short-lived NRF2^WT^ (half-life= ∼20min ^34^), we examined the effects of PYR and MIT on NRF2 half-life. CHX chase experiments showed that PYR (10 µM) and MIT (20 nM), but not MTX (50 nM), significantly shortened NRF2 half-life in *NRF2*^*W24C*^-KYSE70 from 66.7 min to 34.9 min and 42.7 min, respectively (Figure 3A). Similarly, PYR (10 µM) and MIT (20 nM) shortened NRF2 half-life in *NRF2*^*D77V*^-KYSE180 cells (Figure 3B), but not in *NRF2*^*WT*^-KYSE450 cells (Figure 3C).

**Figure 3.**
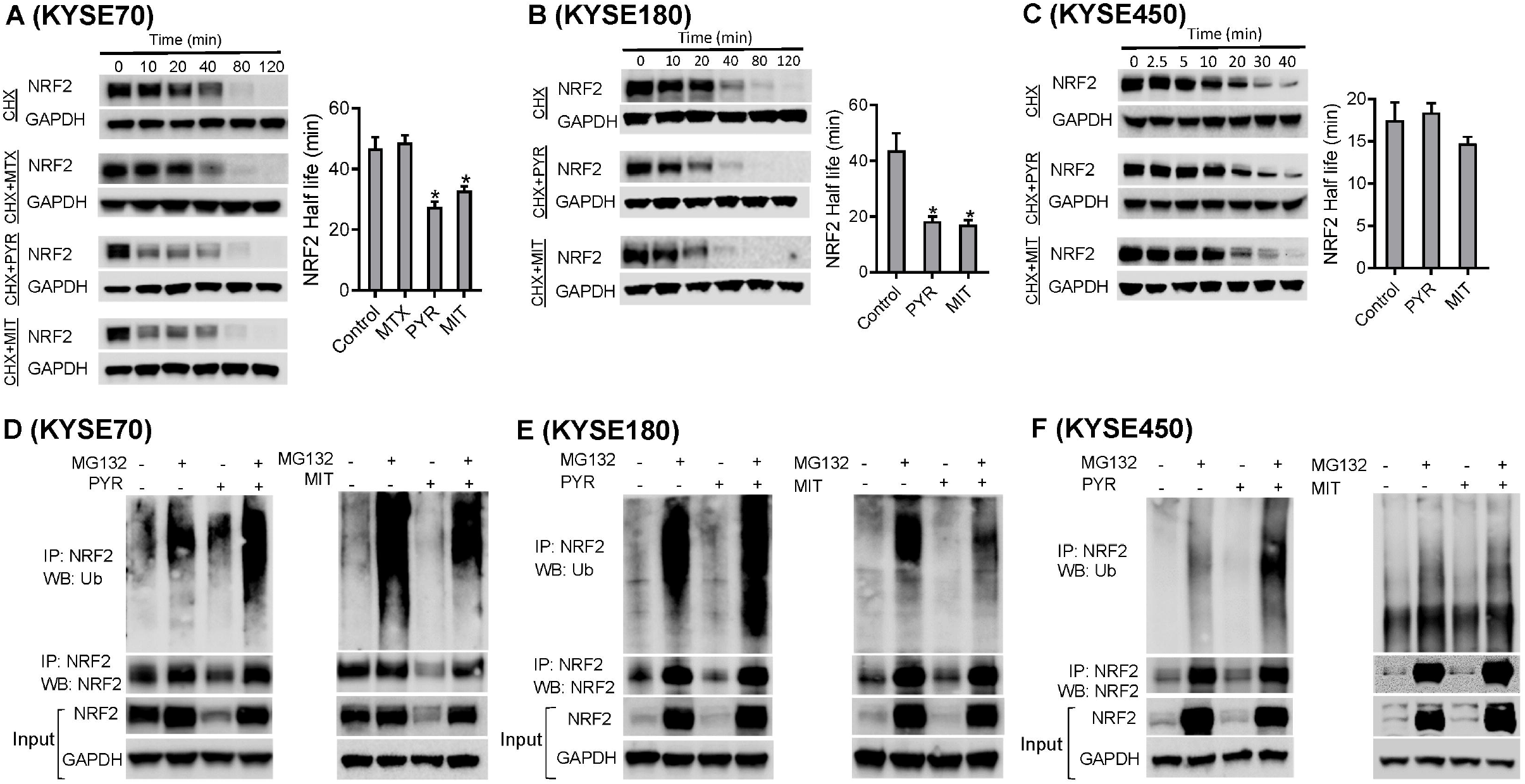
PYR shortened NRF2 half-life and promoted NRF2 ubiquitination in ESCC cells. (A) NRF2 half-life was shortened by PYR (10 µM) and MIT (20 nM), but not by MTX (50 nM), in *NRF2*^*W24C*^-KYSE70 cells. (B) NRF2 half-life was reduced by PYR (10 µM) and MIT (20 nM) in *NRF2*^*D77V*^-KYSE180 cells. (C) NRF2 half-life was not changed by PYR (10 µM) and MIT (20 nM) in KYSE450 cells. (D) PYR (10 µM), but not MIT (20 nM), promoted NRF2 ubiquitination in KYSE70 cells in the presence or absence of MG132 (a proteasomal inhibitor). (E) PYR (10 µM), but not MIT (20 nM), promoted NRF2 ubiquitination in KYSE180 cells in the presence or absence of MG132. (F) PYR (10 µM), but not MIT (20 nM), slightly promoted NRF2 ubiquitination in KYSE450 cells only in the presence of MG132. * p<0.05

We next investigated the effects of PYR and MIT on NRF2 ubiquitination and degradation. Using NRF2-IP and ubiquitin Western, we found that PYR (10 μM), but not MIT (20 nM), promoted NRF2 ubiquitination in the presence of MG132 (a proteasome inhibitor) in KYSE70, KYSE180, and KYSE450 cells (Figure 3D, E, F). Taken together, our data suggested that PYR may inhibit NRF2 expression in *NRF2*^*Mut*^-ESCC cells by promoting NRF2 ubiquitination and degradation, and thus reducing NRF2 half-life.

### NRF2-inhibitory effects of PYR on the NRF2^high^ esophageal phenotype *in vivo*

We chose to test the *in vivo* efficacy of PYR instead of MIT because PYR has recently been tested in clinical trials on cancer. PYR has a better safety profile than MIT and cytotoxicity of MIT may complicate the analysis. A mouse model was first established by expressing a mutant allele in the mouse esophagus (*Sox2CreER;LSL-Nrf2*^*E79Q/+*^)(Figure 4A). Our previous experiment using the *K14Cre;LSL-Nrf2*^*E79Q/+*^ mice has shown that expression of the mutant allele resulted in squamous epithelial hyperplasia in the tongue, esophagus, and forestomach ^25^. Four weeks after tamoxifen induction, NRF2, NRF2 transcriptional targets (GCLC, GCLM) and a keratinization marker (Loricrin) were overexpressed in the mutant esophagus, as detected by Western blotting and IHC (Figure 4B, C). An increased number of BrdU^+^ cells and increased thickness of the keratinized layer indicated hyperplasia and hyperkeratosis of the esophageal squamous epithelium. At Week 6, mild dysplasia developed (Figure 4C). However, up to 20 weeks after tamoxifen induction, no tumors were found in the mouse esophagus (data not shown), suggesting that NRF2 activation likely drives cancer progression but not cancer initiation. Using ^18^F-FDG PET/CT, we further found that this mutant allele resulted in hyperactive glycolysis (increased ^18^F-FDG uptake) after tamoxifen induction (Figure 4D).

**Figure 4.**
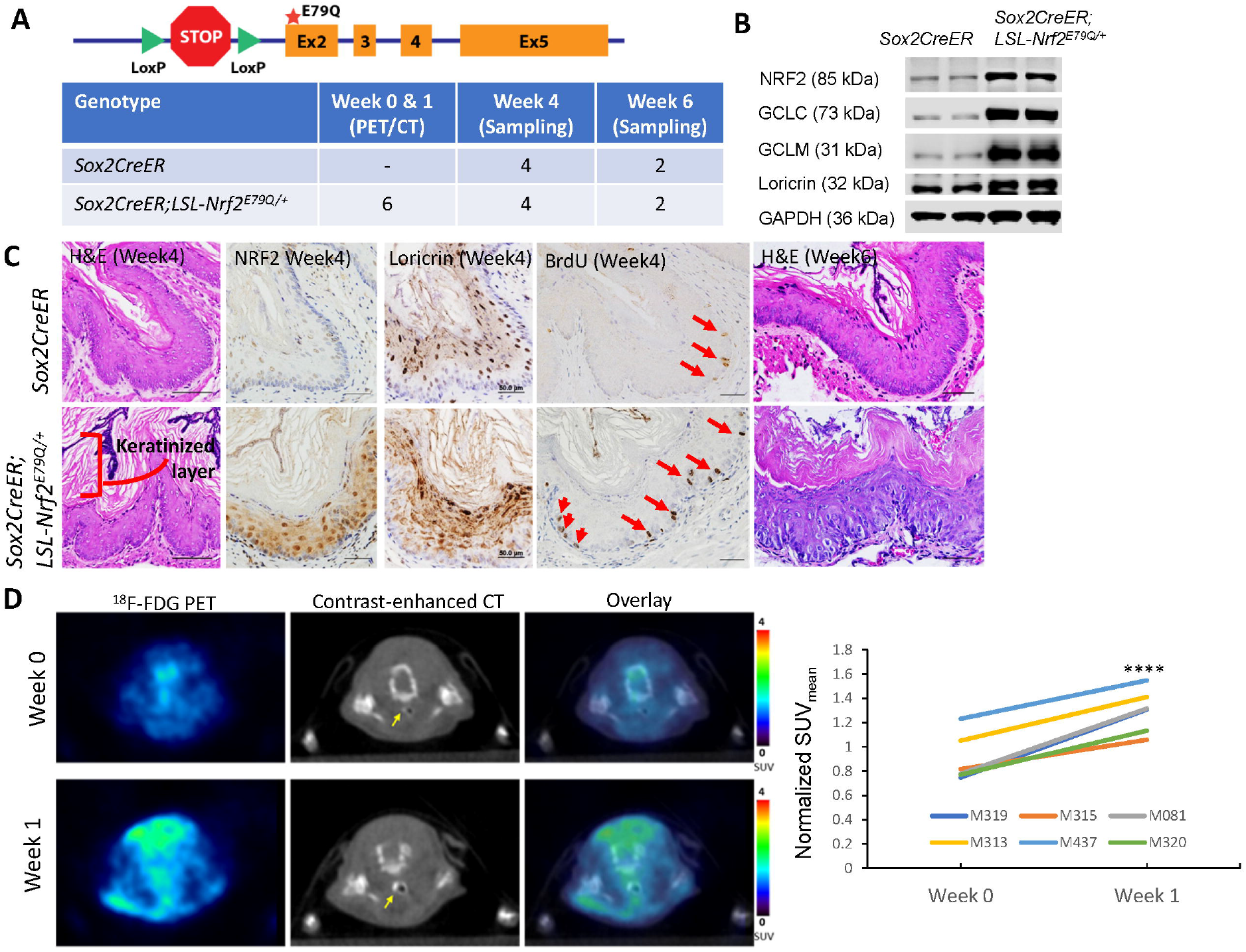
NRF2^high^ esophageal phenotype in Sox2CreER;LSL-Nrf2^E79Q/+^ mice. (A) Schematic figure of the *LSL-Nrf2*^*E79Q*^ allele and experimental design. (B) Expression of NRF2, NRF2 target genes (GCLC and GCLM), and a keratinization marker (loricrin) in the esophageal epithelium of *Sox2CreER;LSL-Nrf2*^*E79Q/+*^ mice as detected with Western blotting; (C) Histology of the esophageal epithelium at 4 and 6 weeks after tamoxifen activation (H&E) and expression of NRF2, loricrin, and a proliferation marker (BrdU) in the mouse esophageal epithelium as detected with IHC. (D) ^18^F-FDG PET/CT of mouse esophagus before tamoxifen induction (Week 0) and after tamoxifen induction (Week 1). Transverse views of the esophagus (arrow, highlighted by the contrast agent) of one mouse (M313) are shown. ****, p<0.001.

Using three doses of PYR (15, 30, and 60 mg/kg/day) in adult wild-type mice, we did not observe any negative impact on the body weight and general health of the mice. However, PYR at the dose of 60 mg/kg/day resulted in myelosuppression including leukopenia and lymphopenia (Table S2). Thus, we chose 30 mg/kg/day *p*.*o*. as the dose for subsequent experiments to investigate the NRF2-inhibitory effects of PYR on the NRF2^high^ esophageal phenotype in our mouse model.

In the first animal experiment, PYR was given together with tamoxifen (Figure 5A). PYR (30mg/kg/day, *p*.*o*., for 5 weeks) reduced the expression of NRF2 and NRF2 target genes (GCLC, GCLM, PKM2) and a cell proliferation marker (BrdU incorporation) which was upregulated by the mutant allele, as detected by western blotting and IHC (Figure 5C). In the second animal experiment, PYR was given after tamoxifen induction (Figure S7A). In the esophageal epithelium, PYR significantly inhibited the expression of NRF2 and its target genes (HMOX1 and AKR1C3). NQO1 expression in the esophageal epithelium was inhibited, yet without statistical significance. However, it remains unclear why one mouse did not respond to PYR treatment. In the forestomach, NRF2 expression was significantly inhibited by PYR as well (Figure S7B).

**Figure 5.**
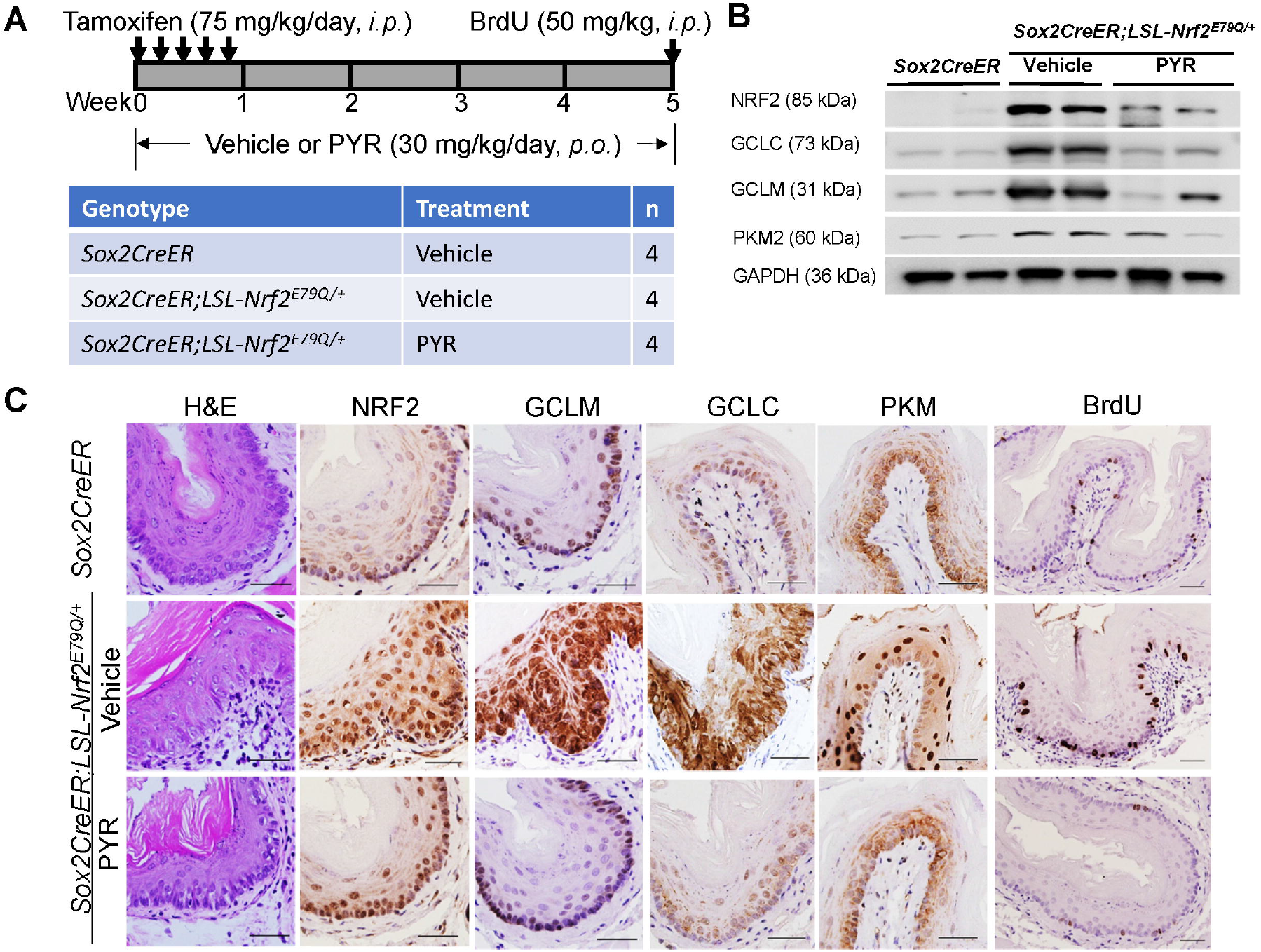
PYR inhibited NRF2 expression in the esophageal epithelium of *Sox2CreER;LSL-Nrf2*^*E79Q/+*^ mice. (A) Experimental design to test the effect of PYR (30mg/kg/day, *p*.*o*.) on the NRF2^high^ phenotype in the mouse esophagus; (B) Expression of NRF2 and its target genes (GCLC, GCLM, PKM2) in the esophageal epithelium of *Sox2CreER;LSL-Nrf2*^*E79Q/+*^ mice as detected with Western blotting; (C) Histology and expression of NRF2, NRF2 target genes, and BrdU in the esophageal epithelium of *Sox2CreER;LSL-Nrf2*^*E79Q/+*^ mice as detected with IHC. Scale bar=50 µm

## Discussion

We identified multiple potential NRF2 inhibitors using high-throughput screening of small molecule compound libraries. Because of its low toxicity in humans and the absence of studies testing efficacy in patients with cancer, we characterized the effects of PYR on ESCC using cell culture and GEMM.

PYR inhibits plasmodial DHFR ^23^ and STAT3 in leukemia cells ^29^, down-regulates mutant Cu/Zn superoxide dismutase (an NRF2 transcriptional target) in the cerebrospinal fluid of amyotrophic lateral sclerosis patients ^35^, and suppresses the growth of various cancer cells ^36-40^. In *NRF2*^*Mut*^-ESCC cells, PYR inhibited NRF2 expression in a dose- and time-dependent manner (Figure 2, S3). RNAseq and qPCR data confirmed the downregulation of NRF2 signaling (Figure S4). We excluded inhibition of STAT3, global translation, and NRF2-ARE binding, as possible mechanisms of action in our experimental system (Figure S5). Instead, we found PYR promoted NRF2 ubiquitination and degradation, and thus shortened its half-life in *NRF2*^*Mut*^-ESCC cells (Figure 3).

In agreement with a previous study^41^, we also identified MIT as a candidate NRF2 inhibitor. Although it did not impact NRF2 ubiquitination, it did shorten NRF2 half-life (Figure 3). MIT is FDA-approved for the treatment of numerous human cancers, but not yet for esophageal cancer ^24^. It has been reported to induce transitory subjective and objective response without significant local or systemic side effects in five patients with inoperable, recurrent esophageal cancer after local administration ^42^. Mechanistically, MIT mainly acts through intercalation with the DNA molecule, which in turn causes single- and double-stranded disruptions and suppresses DNA repair via inhibition of topoisomerase II. It can also intercalate with GC-rich mRNAs (e.g., *HIF1*_α_, *tau* pre-mRNA) ^43, 44^ and interact with proteins (e.g., PIM1 kinase) and lipids non-covalently ^24^. Since the promoter of *NFE2L2* mRNA is very GC-rich ^45^, interference with *NFE2L2* mRNA translation may be a mechanism of action. C_max_ of MIT in the plasma reached as high as 6.43±2.43 mg/L (12.43±4.70 µM) at the dose of 90 mg/m^2^ after intravenous injection. Plasma concentration of MIT remained as high as 4.65±2.35 µg/L (8.99±4.54 nM) at 96 h after injection ^46^. Therefore, the concentrations used in this study are clinically achievable. Further studies are warranted to elucidate MIT’s mechanisms of action and its NRF2-inhibitory effect *in vivo*.

Five strategies have been proposed to target the NRF2 signaling pathway for cancer therapy: (1) transcriptional downregulation of *NRF2*; (2) increased degradation of *NRF2* mRNA or decreased translation; (3) enhancement of NRF2 degradation through up-regulation/activation of KEAP1-CUL3, β-TrCP-SCF or HRD1; (4) blocking the dimerization of NRF2 with small MAF proteins; and (5) blocking the NRF2-sMAF DNA-binding domain^47^. Our data suggest that small molecules (e.g., PYR) may inhibit NRF2 by promoting its ubiquitination and degradation. Consistent with our data, PYR has been reported to promote ubiquitination and degradation of AIMP2-DX2 in lung cancer cells ^48^. These data suggest that studies on the NRF2 protein degradation mechanism of PYR may further elucidate its mechanism of action. On the other hand, antimetabolites (e.g., MTX) have been reported to inhibit NRF2 ^49^. DHFR inhibition was reported as a mechanism of action of PYR on lung cancer cell proliferation ^30^. Thus, PYR may also suppress NRF2 expression by inhibiting DHFR.

The most exciting observation is that PYR effectively inhibited the NRF2^high^ esophageal phenotype in mice. Upon tamoxifen induction for 1 week, *Sox2CreER;LSL-Nrf2*^*E79Q/+*^ mice developed typical NRF2^high^ esophageal phenotype (hyperplasia, hyperkeratosis, and overexpression of NRF2 and its target genes) as reported in *Keap1*^*-/-*^ mice^18^ (Figure 4, 5). Consistent with the phenotype of hyperactive glycolysis in *Keap1*^*-/-*^ esophagus^50^, expression of the *Nrf2*^*E79Q*^ mutant significantly increased ^18^F-FDG uptake in the mouse esophagus (Figure 4D). Cmax of PYR was reported as high as 8.04, 18.5, 35.78, and 28.15 µM when given to mice at the dose of 12.5, 25, 50, and 75 mg/kg, *i*.*p*. ^51^. Thus, 30 mg/kg/day is expected to achieve a concentration of ∼10 µM in mouse tissues. At this dose, PYR was not myelosuppressive (Table S2), yet significantly inhibited the expression of NRF2 and its target genes in the esophageal epithelium and forestomach (Figure 5B, 5C, S6B). In humans, the Cmax of PYR reached 8.28 µM after 50mg/day, *p*.*o*., for 3 weeks ^38^. These data suggest that PYR can be an NRF2 inhibitor with a good safety profile for patients.

In summary, using a small molecule screen we have identified PYR as an NRF2 inhibitor, and further validated it as an inhibitor of NRF2 expression *in vitro* and *Nrf2*^*Mut*^-driven esophageal hyperplasia *in vivo*. The NRF2-inhibitory effect of PYR at a clinically achievable and safe dose suggests that PYR is a promising NRF2 inhibitor that may be rapidly tested in the clinic as a radio- and chemo-sensitizer for human *NRF2*^*Mut*^-ESCC. Further exploration of its mechanisms of action may lead to more potent NRF2 inhibitors for future use.

## Abbreviations

CDDO: 2-cyano-3,12-dioxo-oleana-1,9(11)-dien-28-oic acid
CHX: cycloheximide
CP: canonical pathway gene set
CT: computed tomography
DEG: differentially expressed gene
DHFR: dihydrofolate reductase
ESCC: esophageal squamous cell carcinoma
^18^F-FDG: ^18^F-fluorodeoxyglucose
GFP: green fluorescent protein
GEMM: genetically engineered mouse model
GO: gene ontology gene set
GSA: gene set analysis
IHC: immunohistochemical staining
KB: knowledge-based gene set
KEAP1: Kelch-like ECH-associated protein 1
MIT: mitoxantrone
NRF2/NFE2L2: nuclear factor erythroid 2-related factor 2
PET: positron emission tomography
PYR: pyrimethamine
SUVmean: mean standardized uptake value
TF: transcription factor gene set
YFP: yellow fluorescent protein

## Acknowledgment

This work was supported by NIH R01 CA244236 (to XC and MBM), R01 CA216051 (to MBM and BW), U54 CA156735 Full Project 5 (to XC, KPW, and MBM), U54 MD012392 Research Infrastructure Core (to XC and KPW), and Lineberger Cancer Center Development Award (to XC and HY). UNC BRIC Small Animal Imaging Facility performed ^18^F-FDG PET/CT imaging, and the facility is partially supported by NIH P30 CA016086, and the PET/CT equipment was funded by NIH S10 OD023611. The authors would like to thank Ms. Ginger Smith (BRITE) for technical assistance and acknowledge additional support from the Golden LEAF Foundation and the BIOIMPACT Initiative of the State of North Carolina (KPW).

## Author’s contributions

Conceptualization: XC, MBM, KPW, BW

Methodology: CP, ZX, DL, YL, BB, JC, CH, EWL, JEF, JL, KM, VC, YL, BB, HY

Funding acquisition: XC, MBM, KPW, HY

Writing original draft: CP, ZX, XC

Writing, review & editing: MBM, KPW, HY, BW

**Supplementary Table S1.** Antibody information.

| Antigen | Sample | IHC | WB | Catalog No. | Host | mAb/pAb | Company | City, State |
| --- | --- | --- | --- | --- | --- | --- | --- | --- |
| ABCG2 | Human cell | - | 1:500 | sc-69989 | Mouse | mAb | Santa Cruz | Dallas, TX |
| AKR1C3 | Human cell |  | 1:1,000 | PIPA5106891 | Rabbit | pAb | Thermofisher | Waltham, MA |
| BrdU | Mouse tissue | 1:500 | - | ab6326 | Rat | mAb | Abcam | Waltham, MA |
| GAPDH | Mouse tissue | - | 1:20,000 | ab8245 | Mouse | mAb | Abcam | Waltham, MA |
|  | Human cell |  |  |  |  |  |  |  |
| GCLC | Mouse tissue | - | 1:4,000 | ab41463 | Rabbit | mAb | Abcam | Waltham, MA |
|  | Mouse tissue | 1:100 | - | sc-166356 | Mouse | mAb | Santa Cruz | Dallas, TX |
| GCLM | Mouse tissue | 1:100 | 1:1,000 | ab126704 | Rabbit | mAb | Abcam | Waltham, MA |
| HMOX1 | Human cell |  | 1:2,000 | ab52947 | Rabbit | mAb | Abcam | Waltham, MA |
| LAMIN B1 | Human cell | - | 1:2,000 | 12586 | Rabbit | mAb | Cell Signaling | Danvers, MA |
| Loricrin | Mouse tissue | 1:500 | 1:1,000 | ab85679 | Rabbit | pAb | Abcam | Waltham, MA |
| NQO1 | Human cell | - | 1:10,000 | ab28947 | Mouse | mAb | Abcam | Waltham, MA |
|  | Mouse tissue |  |  |  |  |  |  |  |
| NRF2 | Human cell | - | 1:1,000 | ab62352 | Rabbit | mAb | Abcam | Waltham, MA |
|  | Mouse tissue | 1:50 | - | ab31163 | Rabbit | pAb | Abcam | Waltham, MA |
| PKM2 | Mouse tissue | 1:400 | 1:1,000 | 4053s | Rabbit | mAb | Cell Signaling | Danvers, MA |
| STAT3 | Human cell | - | 1:1,000 | 9139 | Mouse | mAb | Cell Signaling | Danvers, MA |
| pSTAT3 | Human cell | - | 1:2,000 | sc-8059 | Mouse | mAb | Santa Cruz | Dallas, TX |
| TALDO1 | Mouse tissue | 1:25 | 1:1,000 | ab137629 | Rabbit | pAb | Abcam | Waltham, MA |
| TOP2A | Human cell | - | 1:1,000 | LS-B13281 | Mouse | mAb | Lifespan Biosci | Seattle, WA |
| Ubiquitin | Human cell | - | 1:1,000 | sc-53509 | Mouse | mAb | Santa Cruz | Dallas, TX |

**Supplementary Table S2.** Mouse complete blood count of the dose-finding experiment.

| Parameter | Vehicle | 15mg/kg/day | 30mg/kg/day | 60mg/kg/day | P-value<br>(ANOVA) |
| --- | --- | --- | --- | --- | --- |
| Neutrophil (%) | 12.92±3.65 | 15.60±1.65 | 12.48±0.54 | 16.23±2.25 | 0.076647 |
| Neutrophil (/μL) | 1355.00±419.36 | 1831.00±408.76 | 1389.40±214.14 | 1152.75±84.84 | 0.053253 |
| Reticulocyte (%) | 3.42±0.53 | 3.83±0.51 | 4.12±0.44 | 4.03±0.46 | 0.160069 |
| White blood cell (K/μL) | 10.48±1.55 | 11.75±2.49 | 11.10±1.35 | 7.18±0.81 | 0.006249 |
| Absolute Reticulocyte (K/μL) | 370.40±43.60 | 391.75±42.59 | 433.40±48.42 | 418.50±61.26 | 0.238741 |
| Red blood cell (M/μL) | 10.87±0.47 | 10.26±0.28 | 10.514±0.10 | 10.37±0.36 | 0.069741 |
| Hemoglobin (g/dL) | 15.56±0.64 | 14.93±0.30 | 15.36±0.15 | 15.40±0.47 | 0.22043 |
| Lymphocyte (/μL) | 8350.80±1317.29 | 8838.50±1882.37 | 8974.60±1146.01 | 5418.75±789.11 | 0.005304 |
| Lymphocytes (%) | 79.76±5.68 | 75.30±2.52 | 80.80±1.05 | 75.33±2.81 | 0.067851 |
| Hematocrit (%) | 52.18±2.47 | 50.28±1.48 | 51.88±0.98 | 52.58±1.77 | 0.310572 |
| Monocyte (/μL) | 574.60±231.96 | 603.25±131.49 | 477.40±58.40 | 453.50±55.30 | 0.380879 |
| Monocytes (%) | 5.40±1.96 | 5.15±0.41 | 4.34±0.63 | 6.35±0.75 | 0.131002 |
| Eosinophil (/μL) | 189.60±35.93 | 468.00±224.31 | 250.20±60.34 | 145.25±23.51 | 0.004672 |
| Eosinophils (%) | 1.82±0.34 | 3.88±1.48 | 2.30±0.69 | 2.03±0.22 | 0.009788 |
| Mean corpuscular vol (fL) | 48.20±1.10 | 49.00±0.00 | 49.60±0.89 | 50.75±0.96 | 0.005182 |
| Basophil (/μL) | 10.80±1.64 | 9.75±6.55 | 8.80±5.07 | 5.00±6.00 | 0.388585 |
| Basophils (%) | 0.10±0.00 | 0.08±0.05 | 0.08±0.04 | 0.08±0.10 | 0.882927 |
| Mean corpuscular hemoglobin (pg) | 14.32±0.11 | 14.53±0.19 | 14.62±0.19 | 14.85±0.10 | 0.001391 |
| Mean corpuscular hemoglobin concentration (g/dL) | 29.84±0.74 | 29.70±0.29 | 29.60±0.66 | 29.33±0.35 | 0.610224 |
| Platelet Count (K/μL) | 1136.20±151.18 | 1201.75±305.96 | 1157.00±94.81 | 1050.00±93.28 | 0.670916 |

## Supplementary Figures

**Figure S1.**
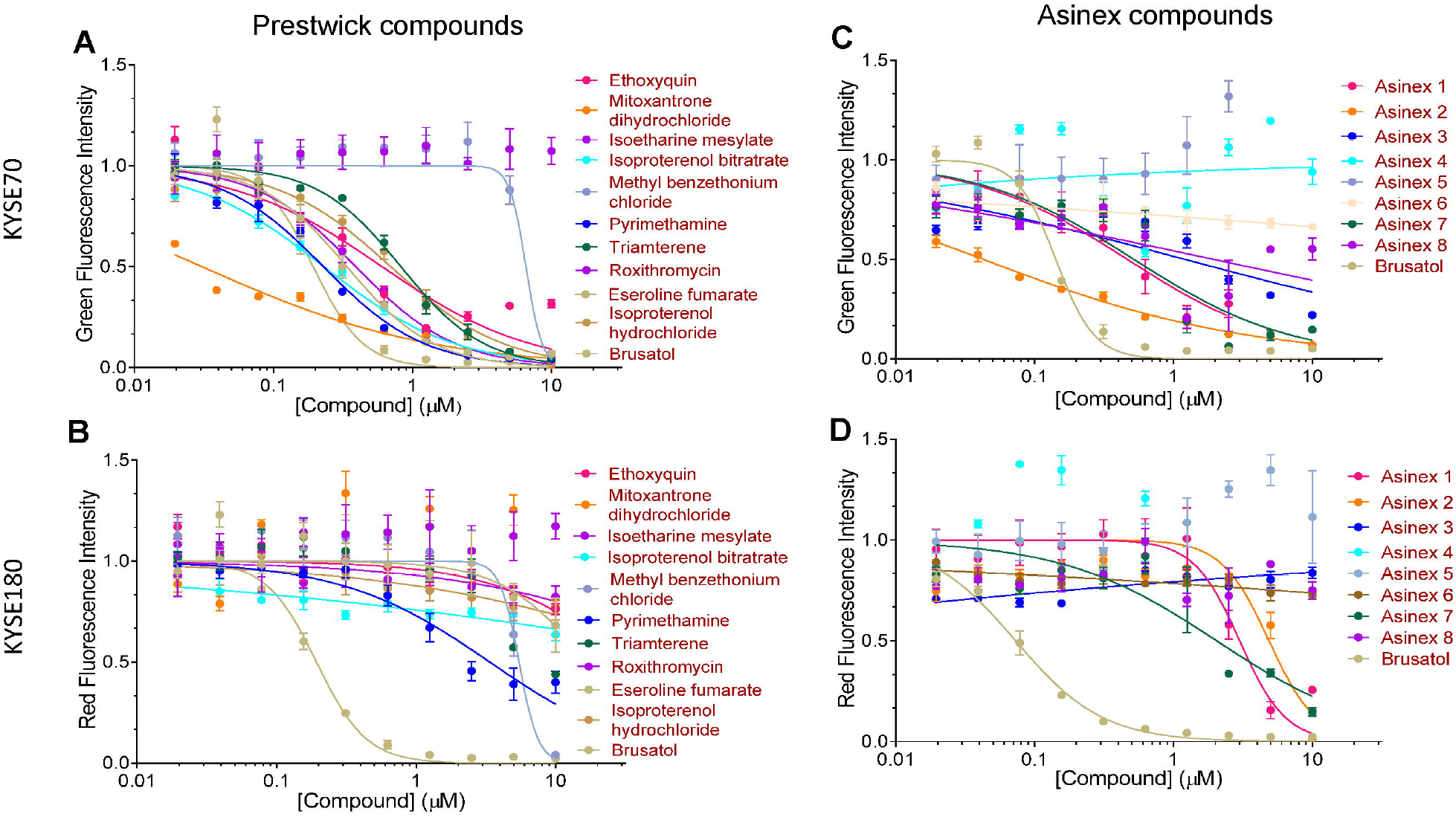
Dose-dependent response of *NQO1-YFP* H1299 cells to hit compounds. (A) Dose-dependent changes of the YPF signal (NRF2 activity) to Prestwick compounds; (B) Dose-dependent changes of the mCherry signal (cell viability) to Prestwick compounds; (C) Dose-dependent changes of the YPF signal (NRF2 activity) to Asinex compounds; (D) Dose-dependent changes of the mCherry signal (cell viability) to Asinex compounds. Brusatol, a known NRF2 inhibitor, served as a positive control.

**Figure S2.**
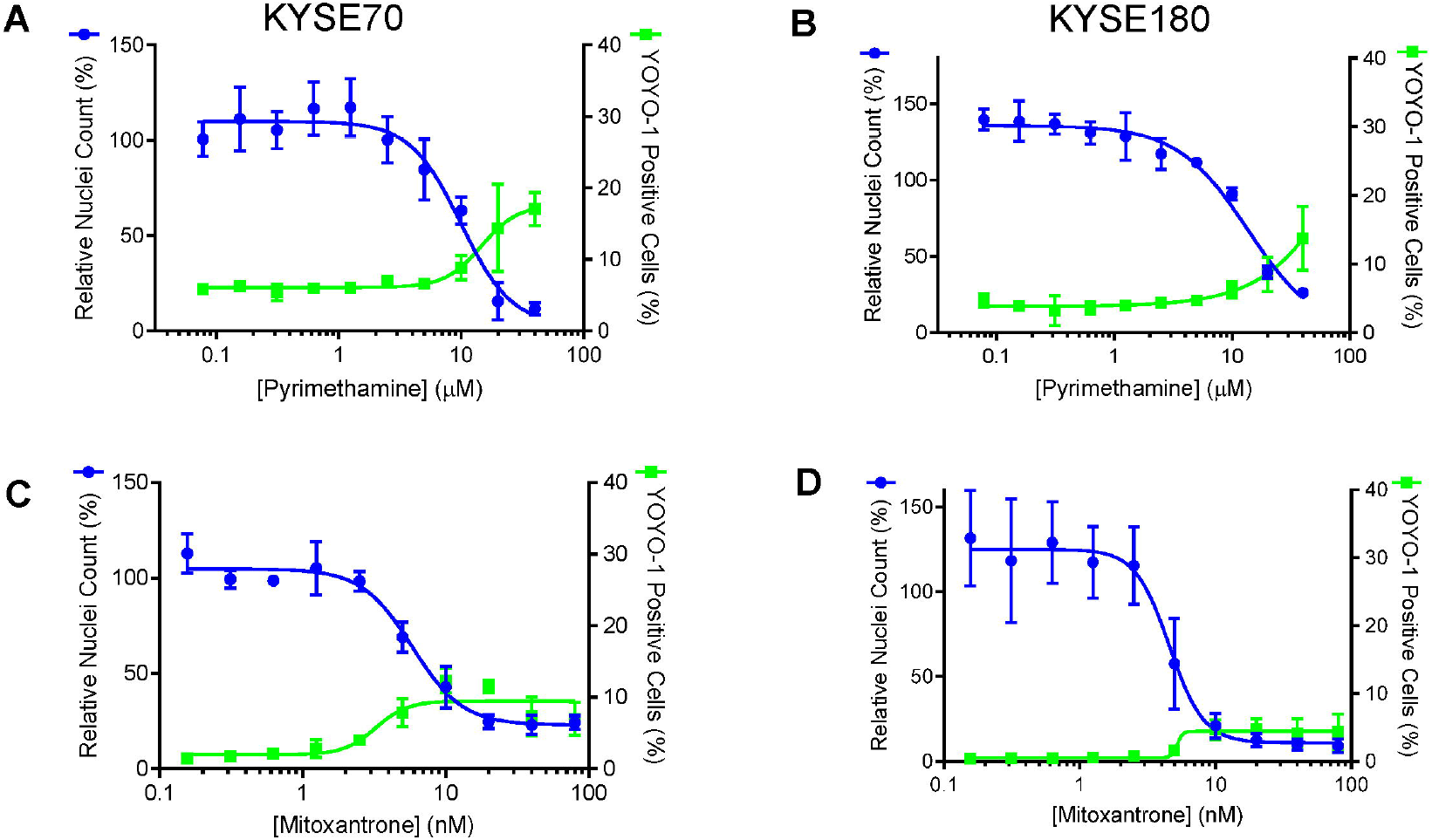
PYR and MIT inhibited cell proliferation in a dose-dependent manner, but does not induce cell death at low concentrations. (A) Dose-dependent effect of PYR on cell proliferation (blue Hoechst signal) and cell death (green YOYO1 signal) in KYSE70 cells; (B) Dose-dependent effect of PYR on cell proliferation (blue Hoechst signal) and cell death (green YOYO1 signal) in KYSE180 cells; (C) Dose-dependent effect of MIT on cell proliferation (blue Hoechst signal) and cell death (green YOYO1 signal) in KYSE70 cells; (D) Dose-dependent effect of MIT on cell proliferation (blue Hoechst signal) and cell death (green YOYO1 signal) in KYSE180 cells.

**Figure S3.**
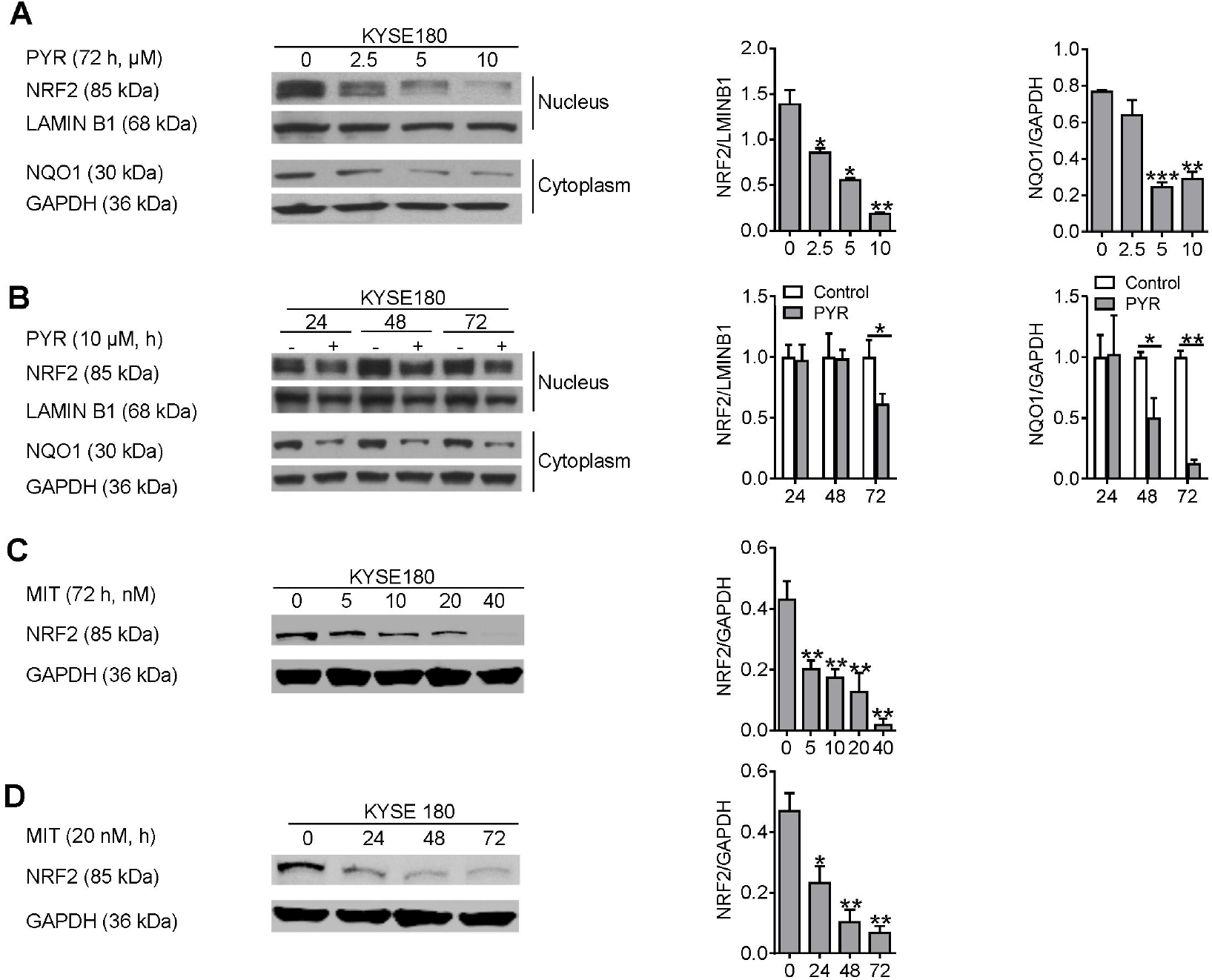
PYR and MIT downregulated NRF2 and NQO1 expression in KYSE180 cells in a dose- and time-dependent manner. (A) Dose-dependent effect of PYR on the expression of nuclear NRF2 and cytoplasmic NQO1 after treatment for 72h; (B) Time-dependent effect of PYR (10 μM) on the expression of nuclear NRF2 and cytoplasmic NQO1; (C) Dose-dependent effect of MIT on NRF2 expression after treatment for 72h; (B) Time-dependent effect of MIT (20 nM) on NRF2 expression.

**Figure S4.**
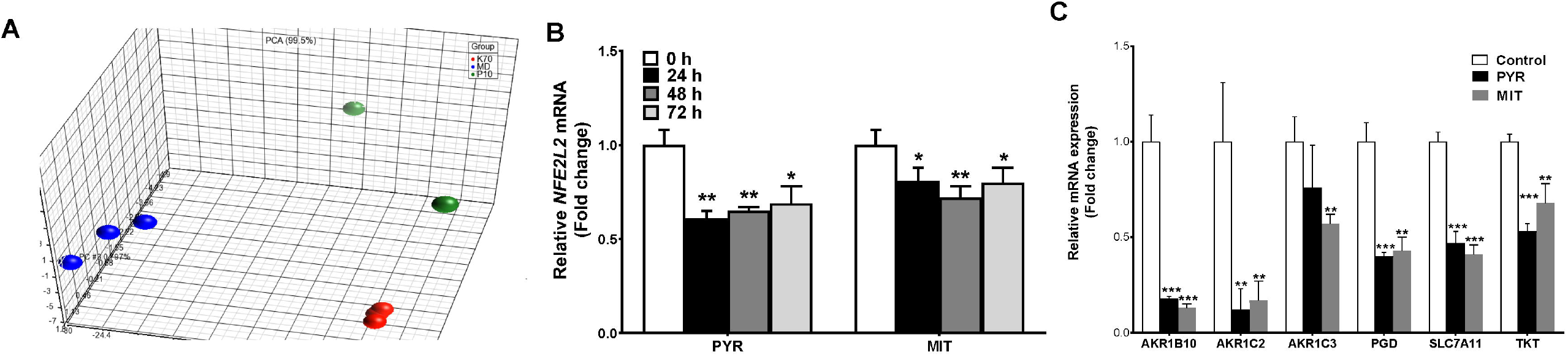
PYR and MIT modulated gene expression in KYSE70 cells. (A) Principle component analysis of RNAseq data of control (green), PYR-treated cells (10 µM for 72 h, red), and MIT-treated cells (20 nM for 72h, blue); (B) Downregulation of *NFE2L2* mRNA by PYR (10 µM for 72 h) and MIT (10 nM for 72 h) as detected by qPCR. (C) Downregulation of NRF2 target genes (*AKR1B10, AKR1C2, AKR1C3, PGD, SLC7A11*, and *TKT*) by PYR and MIT as detected by qPCR.

**Figure S5.**
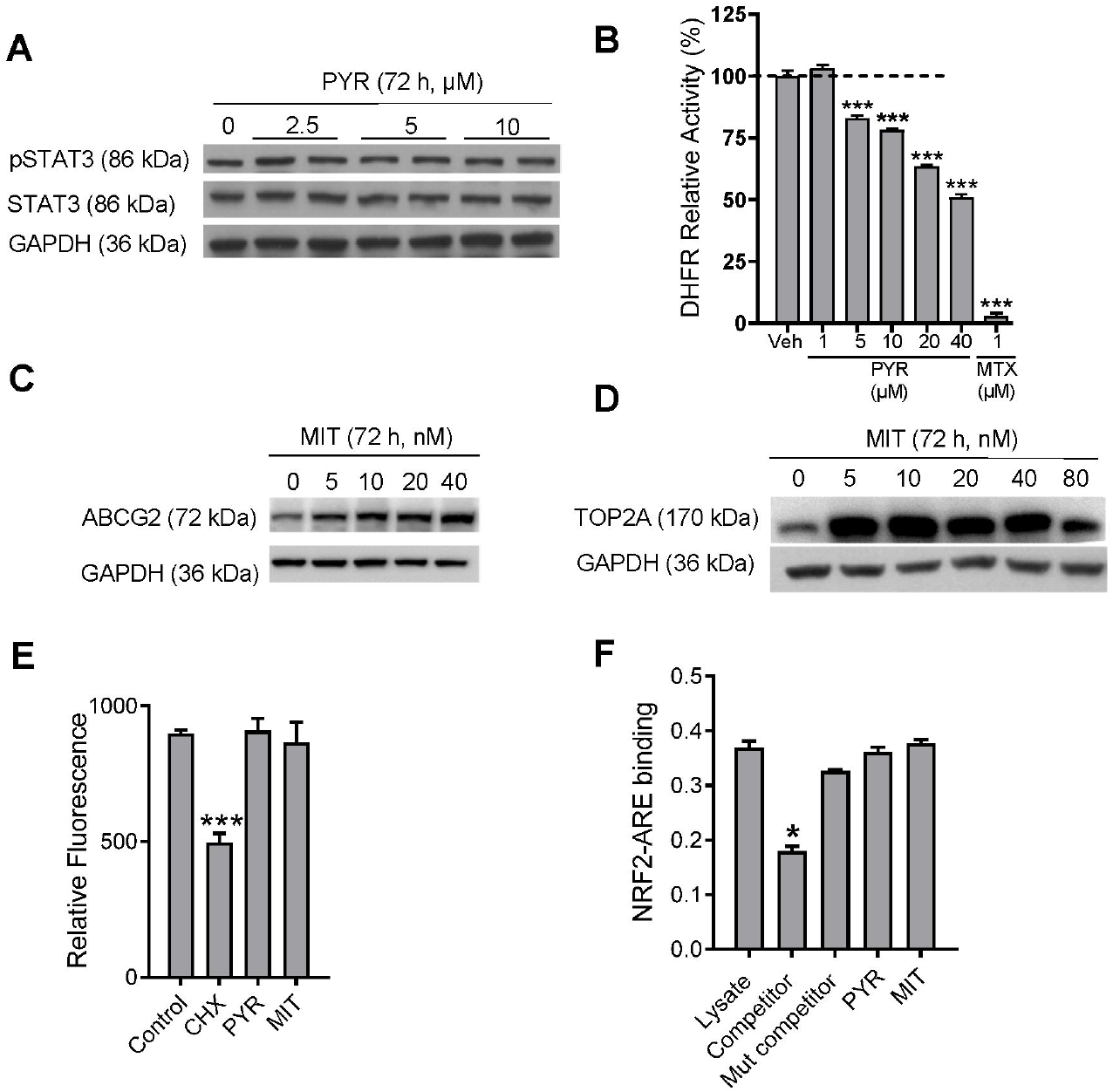
Biochemical effects of PYR and MIT on KYSE70 cells. (A) No significant effect of PYR on STAT3 phosphorylation; (B) Dose-dependent inhibition of DHFR enzymatic activity by PYR with MTX as a control; (C) Increased expression of ABCG2 by MIT; (D) Increase expression of TOP2A by MIT; (E) No significant effects of PYR (10 µM) and MIT (10 nM) on global protein translation with CHX as a control; (F) No significant effects of PYR (10 µM) and MIT (10 nM) on NRF2-ARE binding.

**Figure S6.**
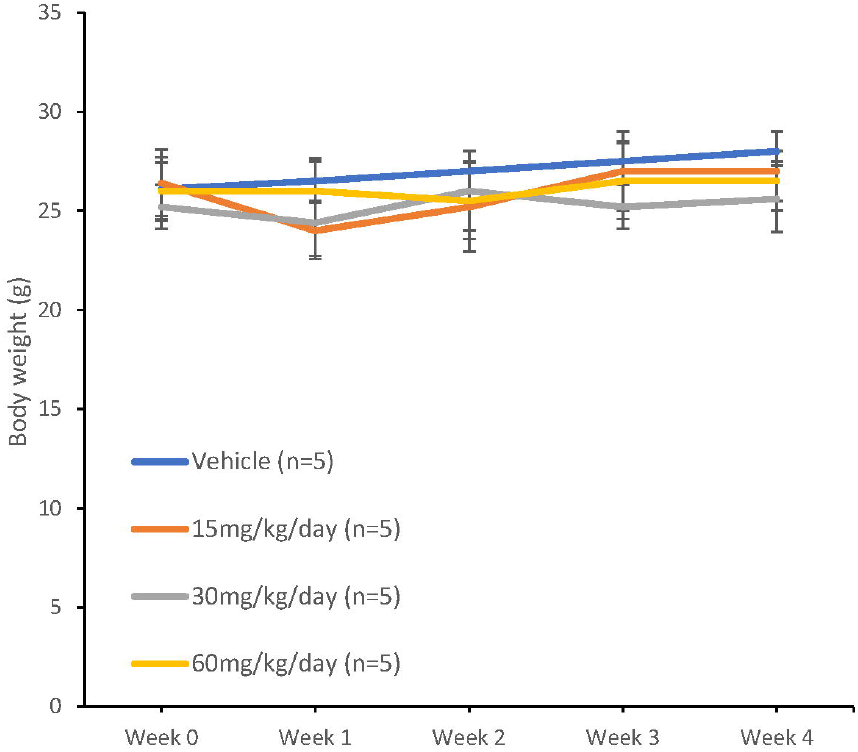
Body weight of wild-type mice (n=5 per group) in the dose-finding experiment.

**Figure S7.**
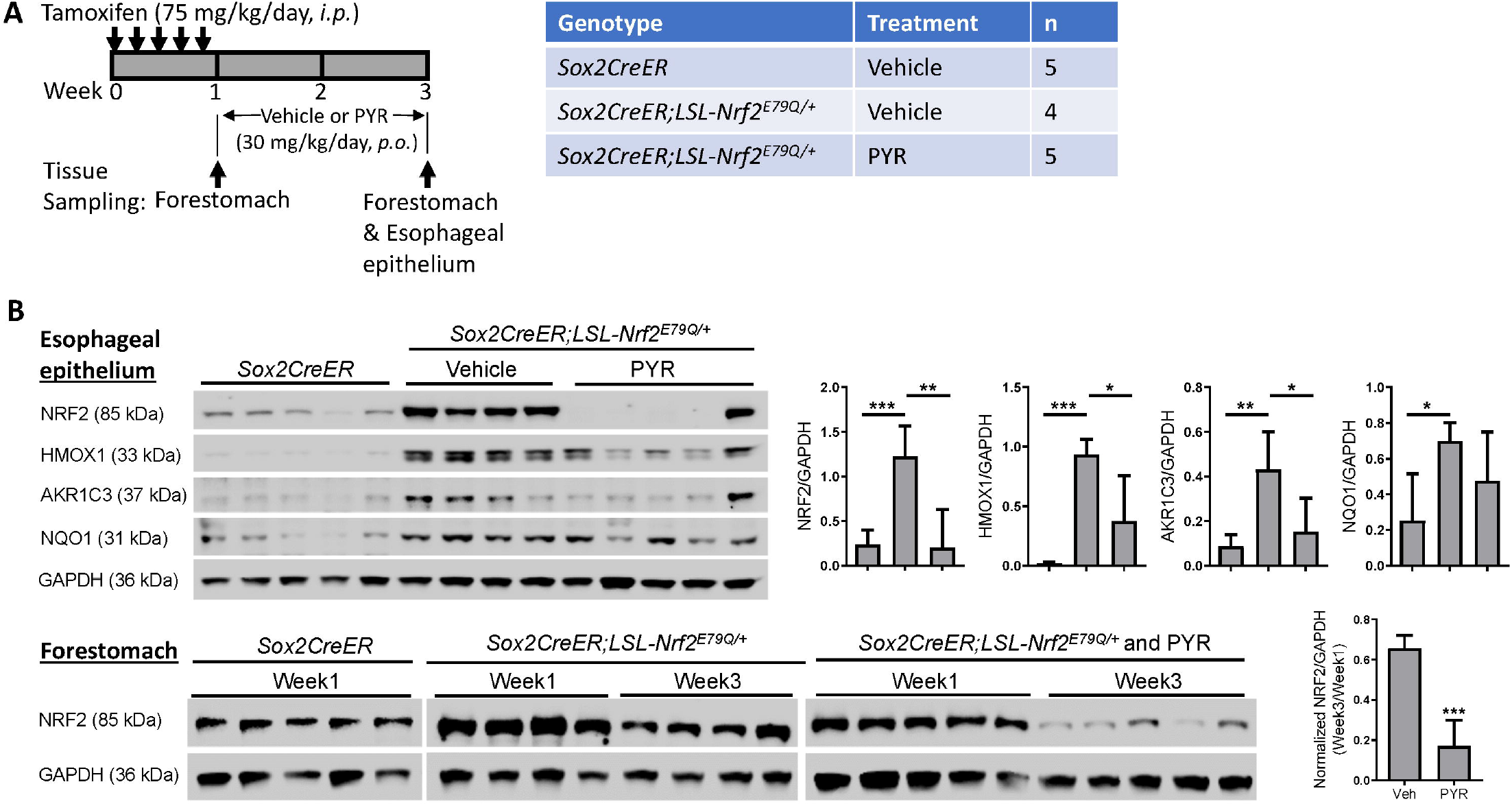
PYR inhibited NRF2 expression in the esophageal epithelium and forestomach of *Sox2CreER;LSL-Nrf2*^*E79Q/+*^ mice after tamoxifen induction. (A) Experimental design to test the effect of PYR (30mg/kg/day, *p*.*o*.) on NRF2 expression in the mouse forestomach and esophagus; (B) Expression of NRF2, HMOX1, AKR1C3, and NQO1 in the esophageal epithelium and forestomach of *Sox2CreER;LSL-Nrf2*^*E79Q/+*^ mice as detected with Western blotting;

## Supplementary Excel file

**Excel S1. Normalized RNAseq data, DEGs, and GSA data in KYSE70 cells: control, PYR (10 µM for 72h), and MIT (20 nM for 72h)**. TF, transcription factor gene set; KB, knowledge-based gene set; GO, gene ontology gene set; CP, canonical pathway gene set.

